# Uncovering the consequences of batch effect associated missing values in omics data analysis

**DOI:** 10.1101/2023.01.30.526187

**Authors:** Harvard Wai Hann Hui, Wilson Wen Bin Goh

## Abstract

Statistical analyses in high-dimensional omics data are often hampered by the presence of batch effects (BEs) and missing values (MVs), but the interaction between these two issues is not well-studied nor understood. MVs may manifest as a BE when their proportions differ across batches. These are termed as Batch-Effect Associated Missing values (BEAMs). We hypothesized that BEAMs in data may introduce bias which can impede the performance of missing value imputation (MVI). To test this, we simulated data with two batches, then introduced over 100 iterations, either 20% and 40% MVs in each batch (BEAMs) or 30% in both (control). K-nearest neighbours (KNN) was then used to perform MVI, in a typical global approach (M1) and a supposed superior batch-sensitized approach (M2). BEs were then corrected using ComBat. The effectiveness of the MVI was evaluated by its imputation accuracy and true and false positive rates. Notably, when BEAMs existed, M2 was generally undesirable as the differing application of MV filtering in M1 and M2 strategies resulted in an overall coverage deficiency. Additionally, both M1 and M2 strategies suffered in the presence of BEAMs, highlighting the need for a novel approach to handle MVI in data with BEAMs.

**Author summary:** Data in high-throughput omics data are often combined from different sources (batches), which creates batch effects in the data. Missing values are a common occurrence in these data, and their proportions are assumed to be equal across batches. However, instances exist when these proportions vary between batches, such as one batch having more missing values than another, resulting in batch effect associated missing values. Missing values are often dealt with through missing value imputation, but whether the variation in missing value proportions across batches affects imputation outcomes is unknown. In this paper, we investigate the consequence of performing imputation when this issue persists. We simulated data with equal and unequal missing value proportions, then assessed the performance of k-nearest neighbours imputation by its imputation accuracy and downstream analysis outcomes. This revealed that unequal missing value proportions worsens imputation and establishes the need for smarter imputation strategies to handle this complication.

## INTRODUCTION

Batch effects (BEs) and missing values (MVs) are common problems in high-throughput omics data, such as genomics and proteomics(1). MVs refer to missing data that should be present in the acquired dataset, and if not properly handled, can negatively impact downstream analyses(2). MVs can cause a loss of statistical power(3), reducing analysis confidence or distorting data distributions, making the data non-representative of the original population(4).

When the proportion of missing values is low, simple omission of missing data may be sufficient. However, when the proportion of missing values is too high to ignore, complete eradication of missing data can result in significant information loss. In these situations, missing value imputation (MVI) techniques can be used to fill in the missing values. MVI techniques are used to estimate missing values based on the available data, which allows the use of powerful analytical techniques such as Principal Component Analysis (PCA) which require complete data. However, it’s important to keep in mind that MVI methods may introduce bias and uncertainty(5), so it’s important to choose the appropriate MVI method for the specific data set and the downstream analysis.

MVs can occur for a variety of reasons, including biological and technical reasons. In some cases, missingness is meaningful and should not be imputed, while in other cases it may be due to limited instrument sensitivity or algorithmic/statistical issues. The mechanism behind MVs is important in determining the appropriate handling procedure(6). MVs can be broadly categorized into three types: Missing not at random (MNAR), missing at random (MAR), and missing completely at random (MCAR)(7). MNAR typically refers to MVs that are missing due to having abundances below the limit of detection (LOD), which is a common type of missingness in omics data(8). While MNAR is explained by the MVs themselves, MAR refers to MVs that can be explained by the observed values. MVs that result from no apparent reason or pattern, and are independent of any unobserved or observed values, are known as MCAR(9). In most omics data, MAR and MCAR are indistinguishable, and thus they are usually assumed to be MAR(10).

MVI covers a wide range of techniques. Simple methods involve replacing an MV with a constant or an estimated value. More complex methods include using statistical correlations or machine learning to transfer information from available data to estimate the missing values. These methods can range from simple mean or median imputation to more sophisticated techniques such as multiple imputation, hot deck imputation, expectation-maximization algorithm, and various machine learning models such as k-nearest neighbors, Random Forest, and neural networks. The choice of the MVI technique will depend on the characteristics of the data, the proportion of missing values, the type of missingness and the downstream analysis(6).

Even though MVI techniques exist, MVs are still considered a problem for several reasons. Determining the specific types and combinations of MVs that plague the data is difficult, making it hard to select the appropriate MVI technique(8). Most omics data are a mixture of MNAR and MAR which makes the selection process even more complex(11). Additionally, the MV filtering threshold, which refers to the proportion of MVs per feature that is acceptable, varies and can range from 5% to 20%(12). This is an important process conducted before MVI that can influence the outcome of downstream analyses. A strict threshold would retain data of higher confidence, whilst losing coverage. A lenient threshold will have greater coverage, but analyses would be more prone to biases. Choosing the appropriate threshold is also a challenging task. Some researchers may choose to avoid MVI as much as possible and instead opt for methods that do not require imputation(13). Ultimately, the choice of MVI technique will depend on the characteristics of the data, the proportion of missing values, the type of missingness, and the downstream analysis(6).

BEs are another common yet complex problem in high-throughput omics data, similar to MVs(1,14). BEs are technical sources of variance between replicates (batches) and are usually caused by factors such as poor experimental design or the combination of data from different studies without sufficient standardization. BEs act as a confounding factor and can have a negative impact on downstream analysis outcomes(1). To address BEs, batch effect correction algorithms (BECAs) are used to estimate and remove BEs from the data(15). Like MVIs, BECAs also make specific assumptions about the BEs, and the choice of a wrong BECA can lead to poor outcomes. A popular BECA is ComBat, which adjusts for batch effects using an empirical Bayes model(16).

Currently, the interaction between MVs and BEs, which are common occurrences in omics data, is not well-studied or understood. Currently, BECAs are designed to deal only on BEs on observable data. They are not designed to handle BEs involving MVs. We termed these Batch Effect Associated Missing values as BEAMs. More specifically, BEAMs are characterized by differential variation of MV proportions across batches, such that one batch can manifest much more MVs than another (Figure 1). In the absence of strategic MVI on a dataset with BEAMs, the different proportions of observed values may cause the MVI to be driven by the batch of least missingness and reduce MVI or subsequent BECA performance. Therefore, it is important to consider both MVs and BEs when analyzing omics data.

**Figure 1.**
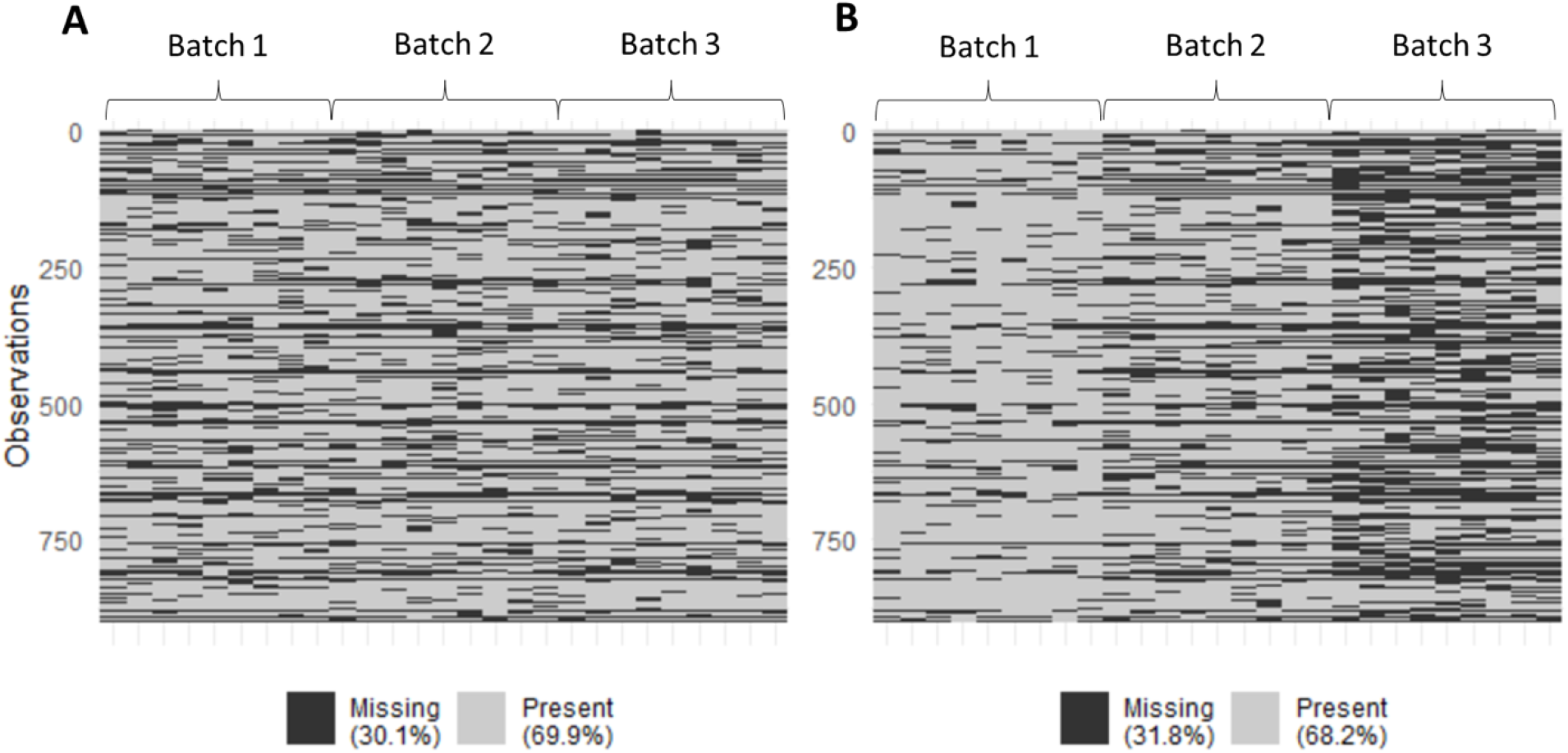
Heatmap visualization of batch effect associated missing values (BEAMs). (A) 30% MVs evenly distributed across batches. (B) 30% MVs disproportionately distributed across batches and are therefore BEAMs.

A previous study evaluated the typical global imputation strategy (termed as M1) against a batch-sensitized imputation approach (termed as M2)(17). It found that an M2 strategy was superior in terms of both imputation accuracy and differential analysis following BE correction. This was due to the retention of pertinent batch information, which improved BECA performances in BE detection and correction. Thus, the general recommendation when performing MVI on data with batches is to employ an M2 approach. However, these tests were conducted on data with evenly distributed MVs, and the utility of M1 and M2 strategies when BEAMs manifest in data is unknown.

In this study, the aim was to investigate the consequences of BEAMs on data when performing MVI using both M1 and M2 strategies. To accomplish this, we first simulated data containing class and batch effects with and without BEAMs. Then, MVI was conducted using the K-nearest neighbors (KNN) algorithm, using both global (M1) and batch-sensitized (M2) strategies. The use of simulated data provides a controlled setting where evaluations based on imputation accuracy and differential analysis can be assessed against a “true” reference. Since varying levels of MVs in each batch between the BEAMs and non-BEAMs data may result in different numbers of features after MV filtering, we also conducted analysis based only on features that were common between both levels of missingness to make fair comparisons. The workflow of the simulation studies is shown in Figure 2.

**Figure 2.**
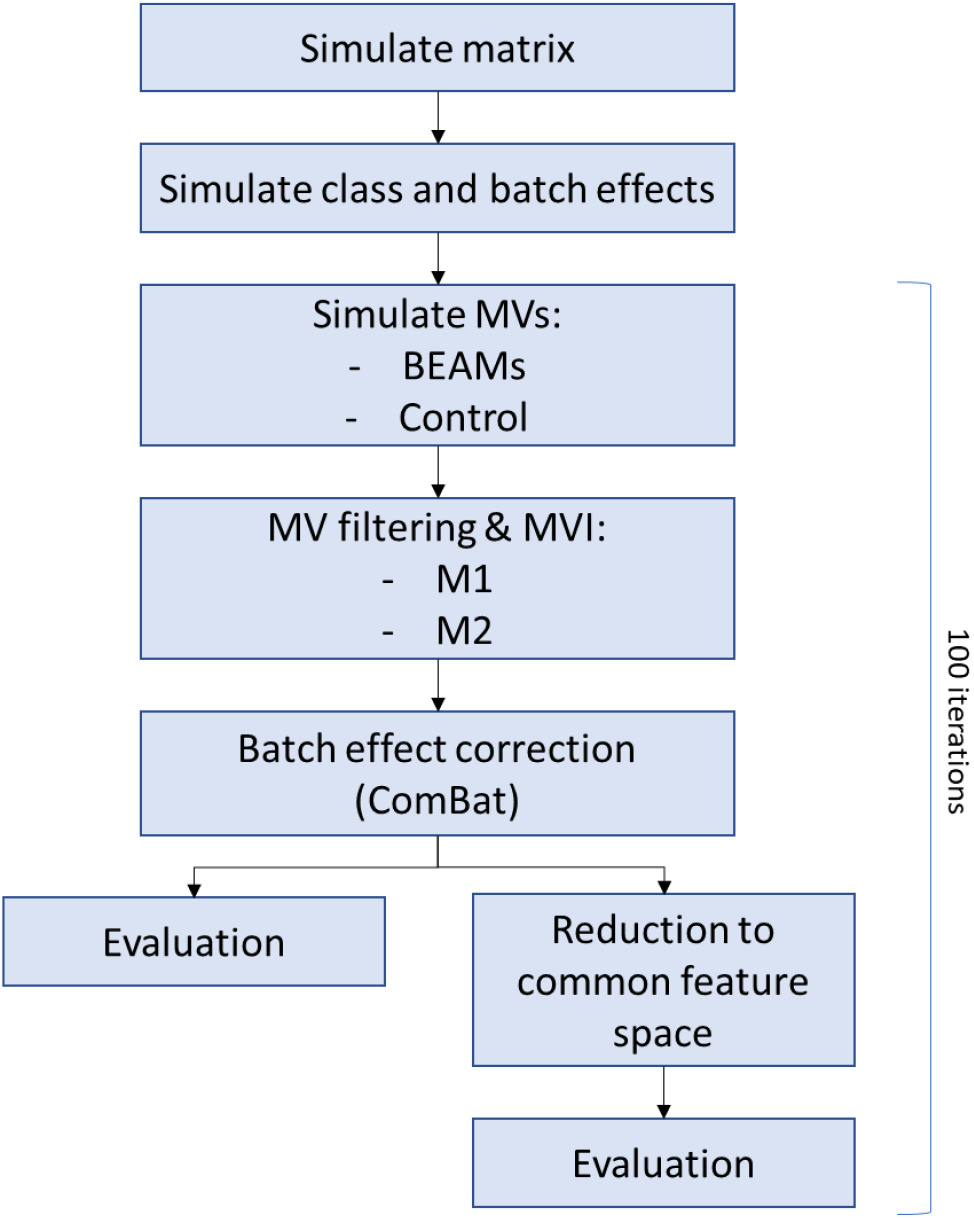
The workflow for simulation studies. 100 simulations were carried out to increase reliability of results.

## Results

### Feature retention post-MV filtering is significantly influenced by BEAMs

As discussed, BEAMs are thought to affect the number of features retained in the data after MV filtering, the process which removes features based on a pre-determined missing threshold. The extent of feature coverage is important as it relates to how much information is retained in our data. Furthermore, certain features could be targets of interest which we want to conserve. Thus, it would be desirable to retain as many features as possible. Although a batch-sensitized (M2) approach is generally considered superior over the standard global (M1) approach(17), we wanted to investigate the differences in feature coverage within these strategies with and without BEAMs, and whether this would affect the selection between M1 and M2.

Thus, we compared the number of features retained with both M1 and M2 approaches in BEAMs and non-BEAMs scenarios. It was found that with moderate BEAMs (20:40%), feature coverage was significantly affected, as an M1 approach retained ~3% more features while an M2 approach retained ~7% less features (Figure 3). These differences widen further with severe BEAMs (10:50%), reaching ~13% more features in M1 and ~17% less features in M2 (Figure S2). With M1, the MV threshold for each feature was set across all batches. This means that when BEAMs are present, the batch with the lowest missingness may aid in fulfilling that threshold criterion for that feature, resulting in greater feature retention with M1. In an M2 approach, the MV threshold was set within each batch. Thus, if a feature failed to meet the threshold for even a single batch, the entire feature would be removed from the data. While this implies that an M2 strategy imputes only on the most confident data, it would be at the expense of an overall loss of coverage.

**Figure 3.**
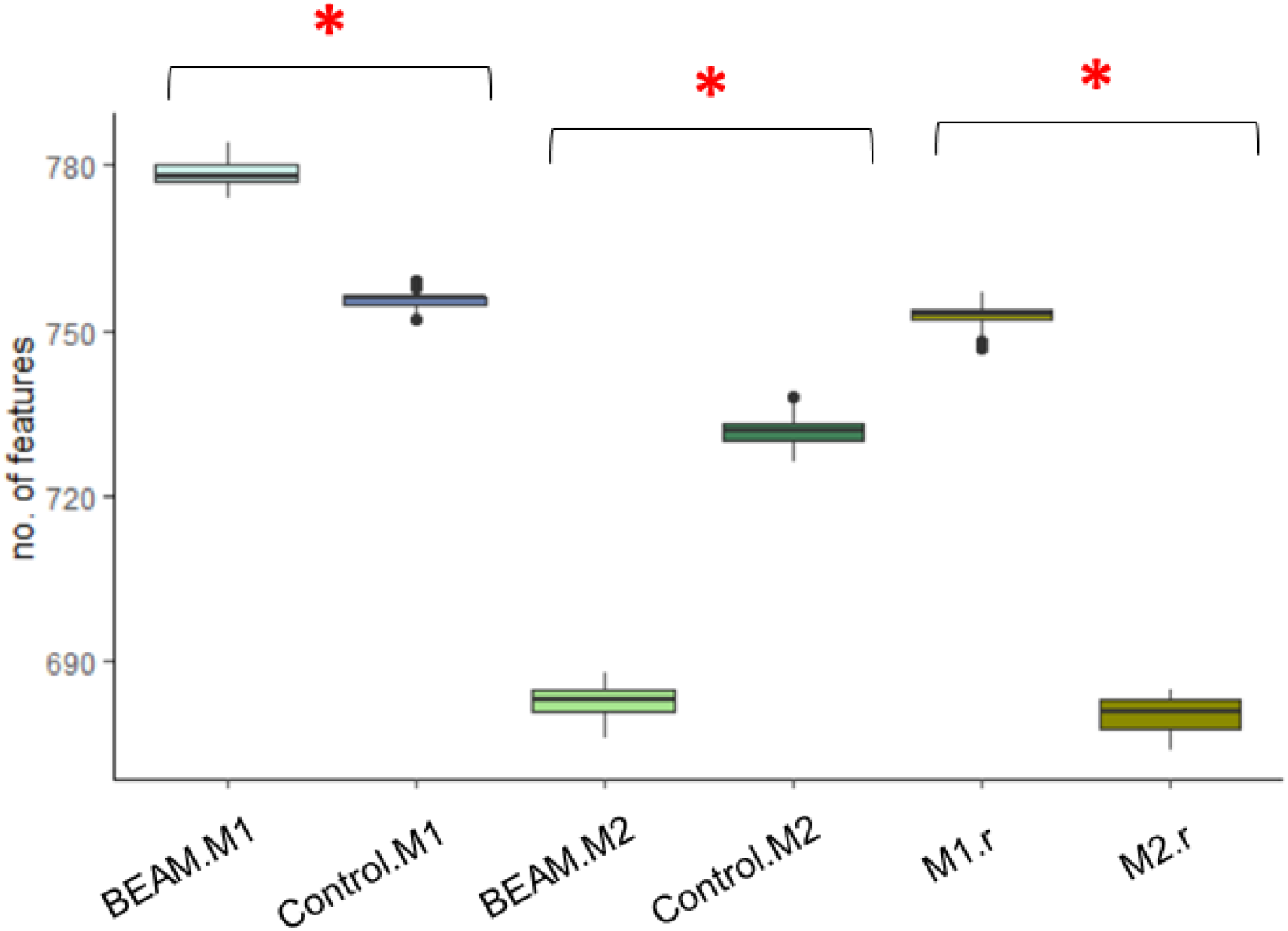
Number of features retained after features with 80% or more MVs were removed in data with moderate BEAMs (20:40% MVs). M1 and M2 with an “r” suffix refer to common features between BEAMs and Control within the same imputation strategy. Asterisks indicate significant difference (p < 0.05).

We then reduced the datasets to the common features retained between the imputed BEAMs and control data in each imputation approach, denoted as “M1.r” and “M2.r” (Figure 3). As expected, most features found in the smaller dataset were found in the larger dataset as well. These reduced datasets were then used to make fairer evaluations between the BEAMs and control imputations. Although comparisons can be made between M1 and M2 with BEAMs in the same feature space, it would not hold much value as the large discrepancy in feature coverage should not be disregarded in the first place.

### BEAMs worsen imputation accuracy regardless of imputation strategy (M1 or M2)

A common strategy to evaluate imputation accuracy is the use of the NRMSE metric, which measures the similarity between the imputed values and the true values. A lower NRMSE value indicates higher similarity and thus greater imputation accuracy.

In the initial comparisons between the BEAMs and Control data, the NRMSE rankings appeared to mirror that of the number of features (Figure 4A). Since this was not a fair comparison, we assessed the BEAMs and Control data within the same imputation strategy reduced to the same features (Figure 4B). The subsequent NRMSE analysis then showed that MVI on data with BEAMs significantly reduced imputation accuracy for these selected features in both M1 and M2 strategies, with a greater extent in M1.

**Figure 4.**
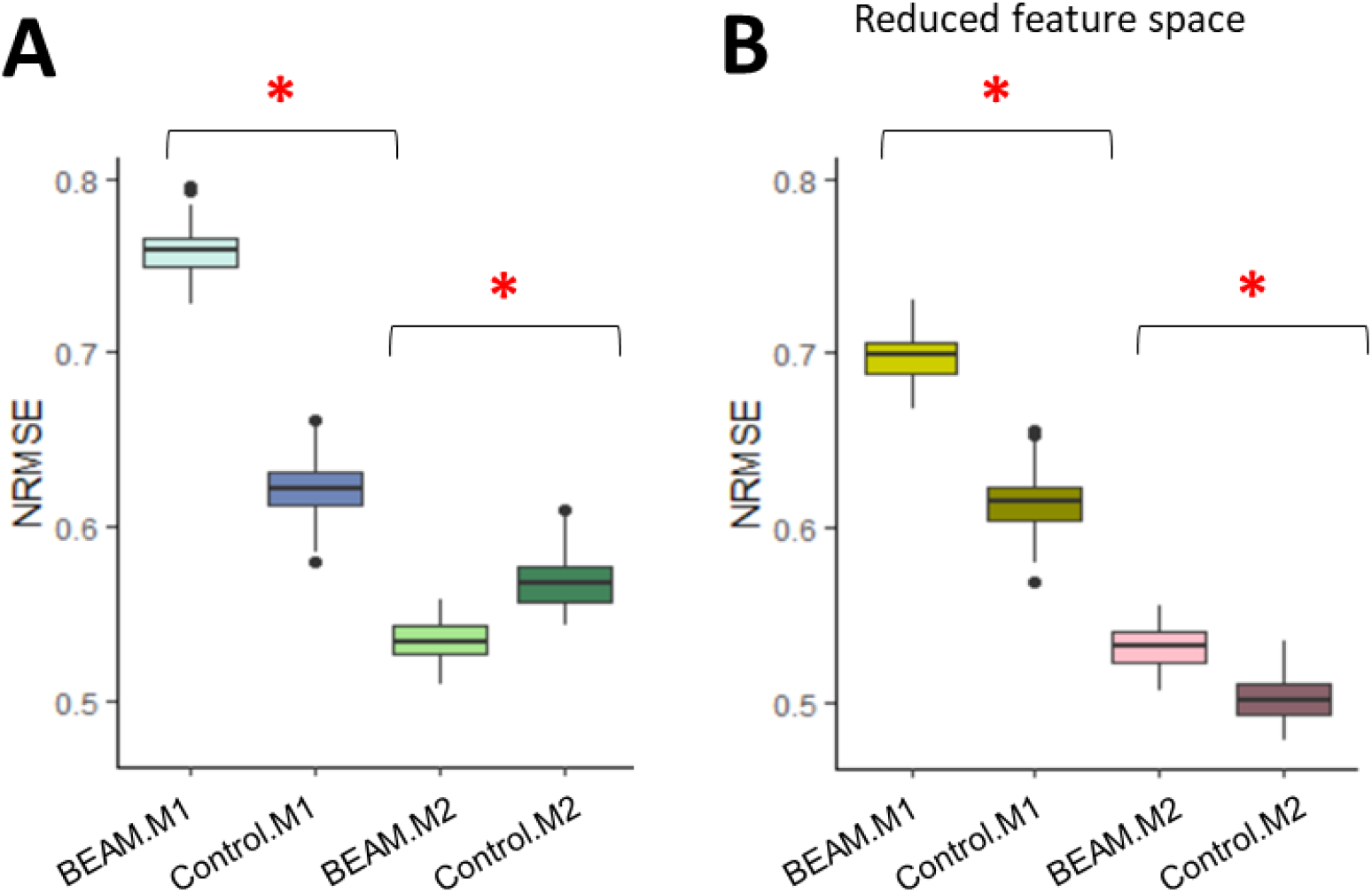
NRMSE outcomes after MVI with and without moderate BEAMs (20:40%). Asterisks indicate significant difference (p < 0.05). (A) Data are analyzed within their own feature space. (B) Data were reduced to the same features within the same imputation strategy (i.e., M1, M2).

In the study, after the 100th iteration, we examined the additional M1 features that were only retained in the BEAMs data. A comparison of the values from the true and imputed datasets before BE correction revealed that the imputation was strongly guided by the batch with lower missingness (Figure 5). This is expected, as when observations are insufficient in one batch, the imputation would inevitably draw information from another batch to form an estimate. Thus, although a BEAMs-infested data may retain more features with an M1 approach, the imputations for these features would be of poor quality and may even mislead analysis.

**Figure 5.**
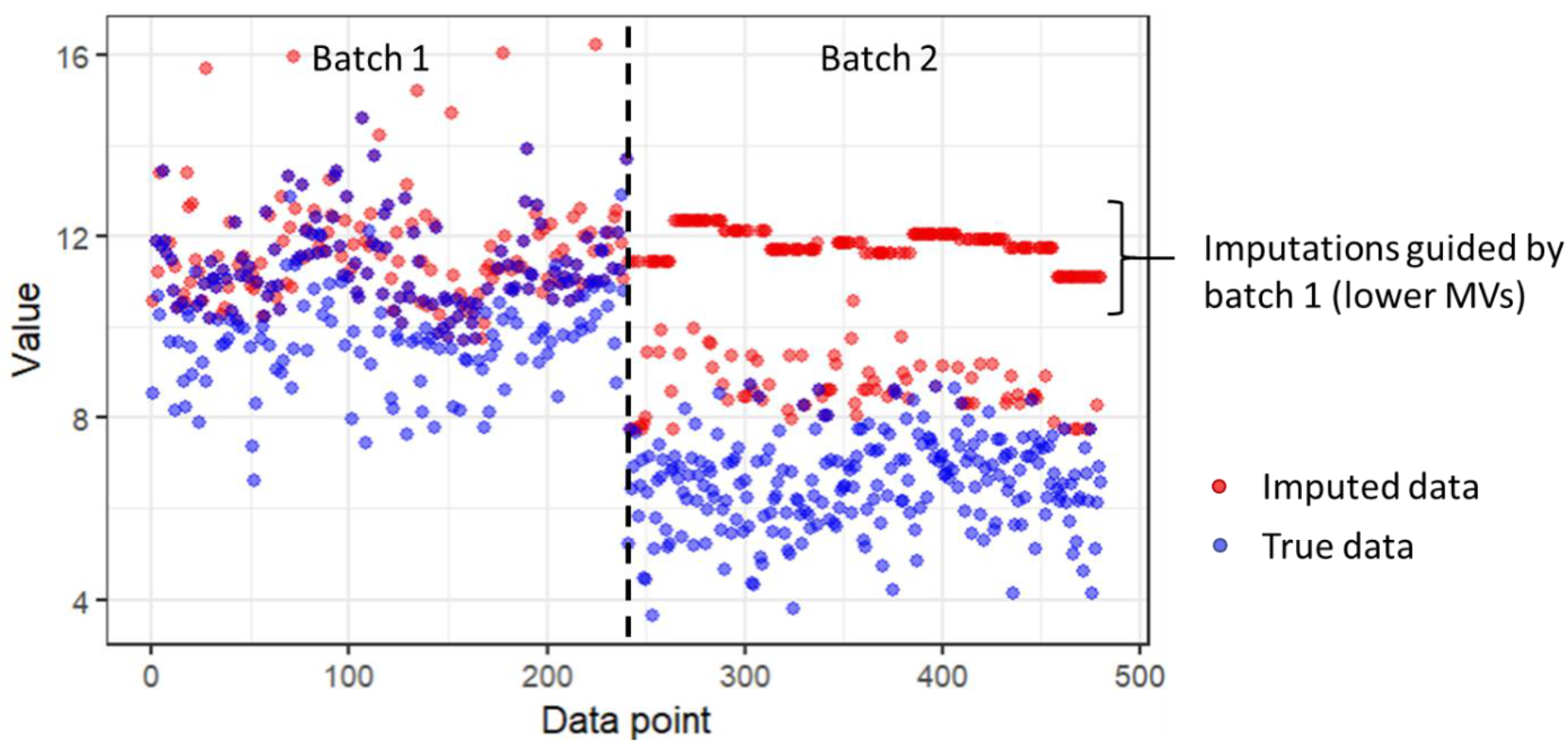
Comparison of values before batch effect correction, in features retained only in M1 with BEAMs data. Red points refer to data points from the imputed data and blue points refer to data points from the original true data. The horizontal intersect separates the data points by their batches.

### Poorer imputation accuracy in BEAMS is generally accompanied by worsened True Positive Rates

After imputation accuracy evaluation, we then determined the extent of which the imputations retained class information, through assessing their TPR and FPR (Figure 6). With moderate BEAMs (20:40% MVs), we found that M1 suffered in TPR (Figure 6A) but not FPR (Figure 6B). M2 with BEAMs without feature reduction showed increased TPR (Figure 6A1), though this was at the expense of an increased FPR as well (Figure 6B1). This, however, might have been a result of the discrepancy in feature space, as these differences were abolished after reduction to the same features (Figures 6A2 & 6B2).

**Figure 6.**
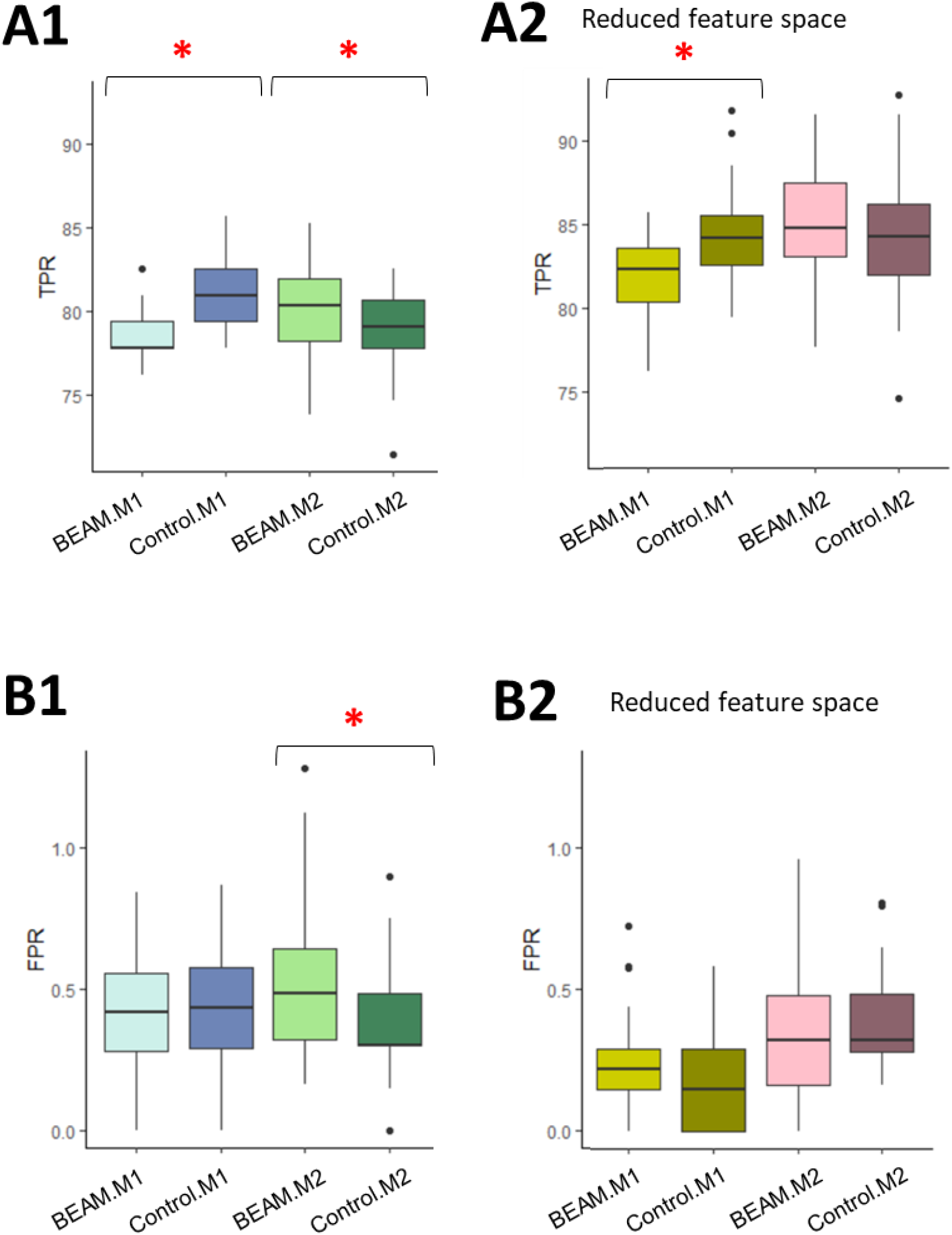
(A) TPR and (B) FPR outcomes after MVI with and without moderate BEAMs (20:40% MVs). Asterisks indicate significant difference (p < 0.05). (1) Data are analyzed within their own feature space. (2) Data were reduced to the same features within the same imputation strategy (i.e., M1, M2).

In a more drastic BEAMs scenario (10:50% MVs), the consequences of MVI on data with BEAMs becomes apparent in both M1 and M2 strategies, with every BEAMs scenario producing worse TPR outcomes than the controls (Figure S4). Whilst the presence of BEAMs did not appear to inflate the FPR with an M2 approach, an M1 approach with BEAMs produced significantly more false positives.

### Sample variance was more closely preserved when MVs are evenly distributed than in BEAMs

To gain a better understanding of why MVI on data with BEAMs was performing worse, we then assessed the intra-sample variance of the dataset from the final iteration, reduced to the same features between BEAMs and non-BEAMs data. Although imperfect, the sample variances obtained in the control data were visibly closer to the original than that of the BEAMs data (Figure 7). In the latter, sample variances appeared to be lower, which might be an indication that the imputation was influenced by a certain batch, causing the variance to shrink.

**Figure 7.**
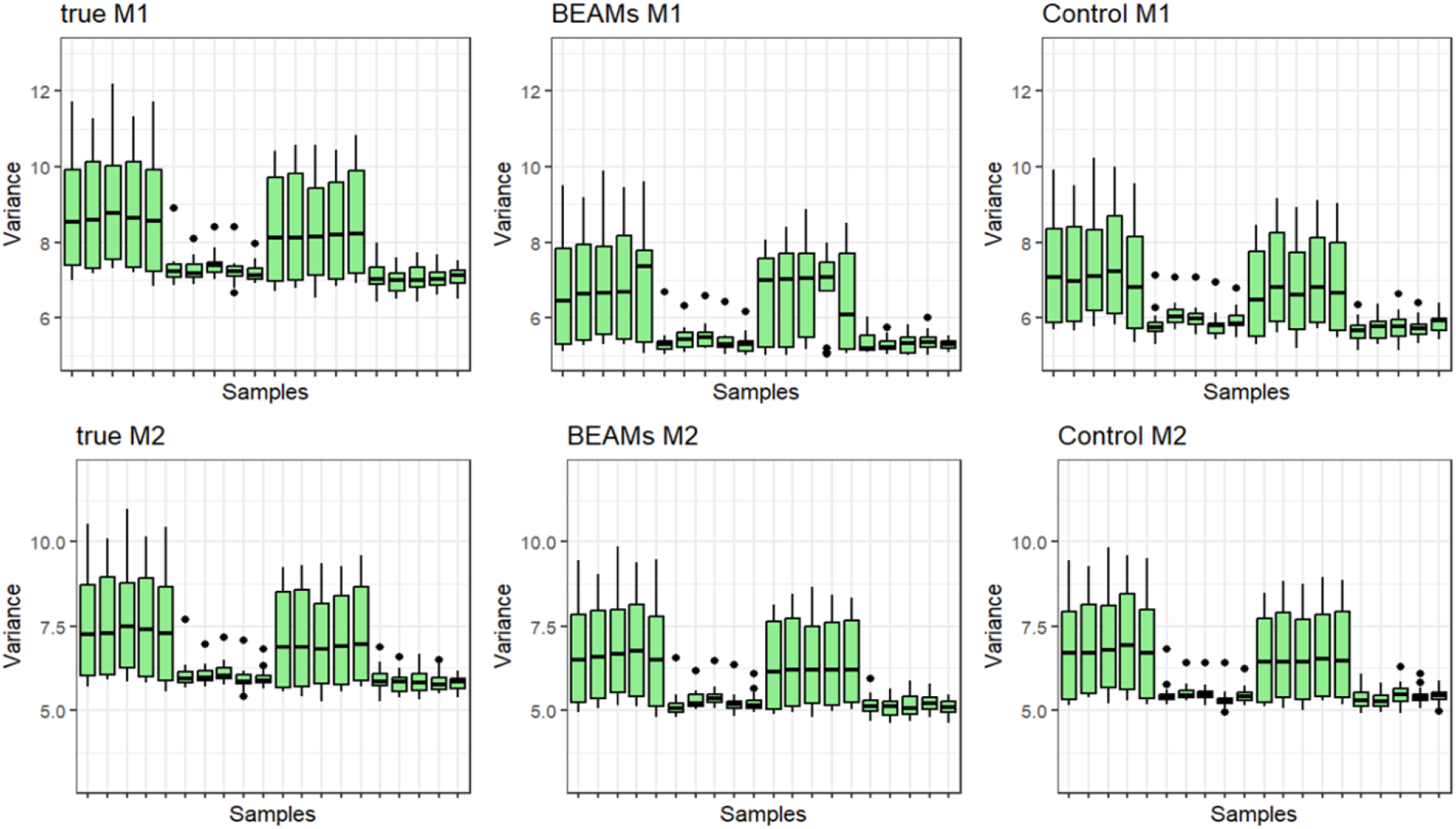
Intra-sample variances of datasets from the final simulation iteration with moderate BEAMs (20:40% MVs). Data were reduced to the same features within the same imputation strategy (i.e., M1, M2).

Earlier, we saw that M2 with moderate BEAMs showed only slight differences compared to the control in the same feature space in TPR and FPR evaluations. When the severity of MV disproportion increased, the consequences of BEAMs became much more apparent. Encouragingly, the sample variances showed consistency with these results, as they were generally similar between the control and moderate BEAMs scenario but deviated from the original when BEAMs were more severe (Figure S5).

### Case study: CPTAC study 6

A real-world example of BEAMs can be seen in the popular CPTAC study 6 dataset(18,19). When data from three machines were combined (therefore forming three batches), we saw that MVs were unevenly distributed across each batch (Figure 8). Specifically, the missing proportions stand around 64%, 33% and 45% in the respective batches, amounting to 47% MVs in total. Thus, this data contains a rather severe case of BEAMs. With real-world BEAMs, we cannot determine the performance of MVI compared to if BEAMs were not present. However, suppose M2 is expected to be the preferred method, determining the number of features retained between M1 and M2 would give an indication as to whether an M2 approach should be considered in the first place.

**Figure 8.**
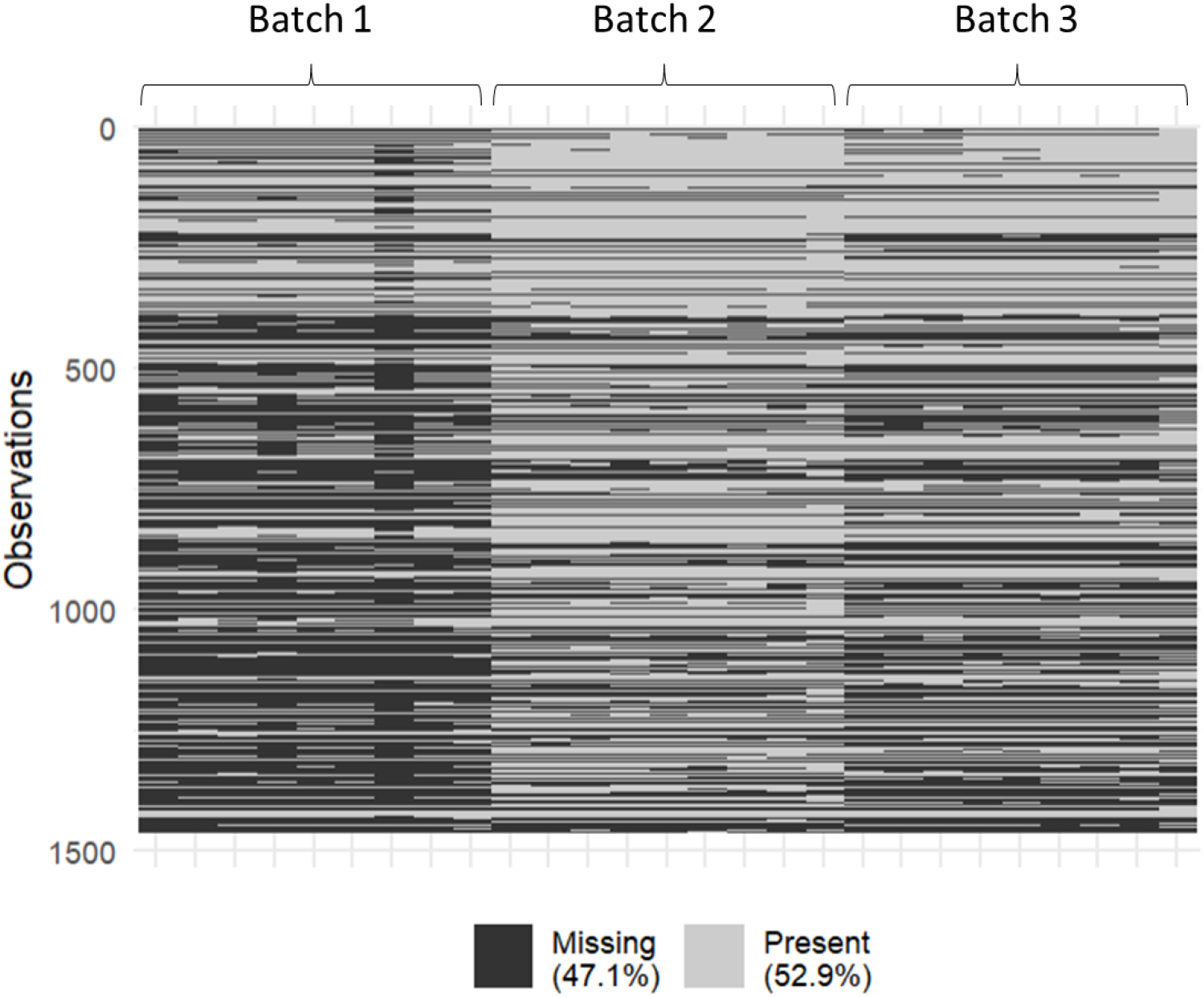
Heatmap of the CPTAC study 6 data from three machines. BEAMs were clearly observed as the machines (batches) displayed clear differences in MV proportions.

Following M1 and M2 MV filtering processes, we find that the resulting data retains only 1024 (30% loss) and 574 (60% loss) features respectively, out of the initial 1462 (Figure 8). In this initial data, 40 spike-ins were the features of interest. M1 retained a total of 30 of these features, whereas M2 retained only 9, demonstrating again the drastic loss of features that can arise with an M2 strategy on data with BEAMs.

## Discussion

### BEAMs worsen KNN MVI regardless of feature retention extent

Post filtering with a predefined MV threshold, the BEAMs data consistently retained more features in M1 and less in M2. Despite having opposing outcomes on M1 and M2 approaches after MV filtering, the negative impact of BEAMs on MVI was clearly seen in both strategies when the same feature space was assessed. This implies that the performance of the MVI was not due to the size of data, but rather, the batch-biased distribution of MVs. Furthermore, after assessing the extra features retained in the BEAMs M1 data, a dilution of batch signal was clearly observed (Figure 5). As the retention of batch signal for BECA detection is important for BE correction, M1 imputation with BEAMs would undoubtedly result in poorer BECA outcomes. Hence, the increased number of features retained in M1 with BEAMs cannot be considered a benefit, as the MVI was not carried out effectively to begin with.

### M2 strategies accommodate moderate cases of BEAMs at the cost of feature retention

In our analyses, we saw that an M2 approach when BEAMs were moderate (20:40%) produced similar performances to when MVs were equally distributed. As mentioned earlier, features that fail to meet the MV threshold in either batch are dropped entirely when using an M2 approach. This naturally reduces the extent of BEAMs after MV filtering, as the batch with more MVs would be more likely to contain features that fail to meet the criterion. However, this does not deal with BEAMs perfectly, as we saw that with severe BEAMs, M2 performance plummets in the same feature space as the control. In any case, when BEAMs are considered moderate, it would not be immediately unwise to use an M2 approach, but we must determine the importance of the features that would potentially be lost in the process. In some situations, the loss of features may skew the data and introduce bias, which can affect downstream analyses outcomes. Furthermore, there is the possibility that a biomarker of interest resides in one of these features but was removed as it contained only enough observations in one batch but not the other, as seen in our CPTAC case study (Figure 9).

**Figure 9.**
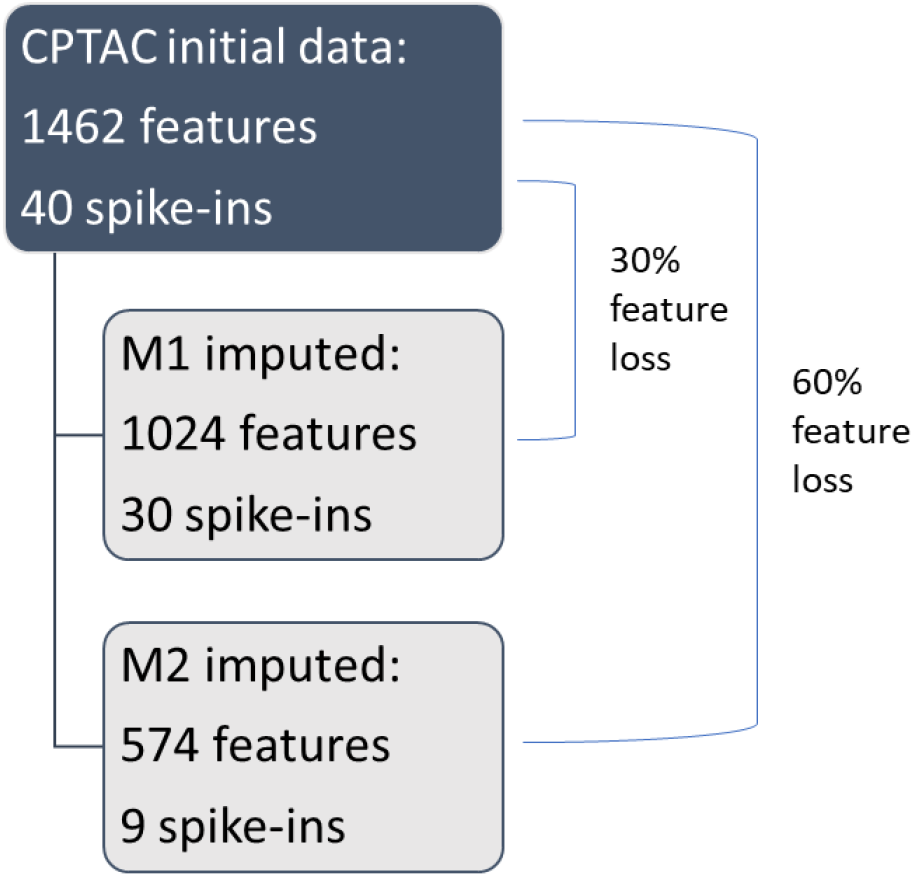
Feature characteristics of the CPTAC study 6 data before and after M1 and M2 MVI.

### Neither M1 nor M2 capable of dealing with BEAMs without drawback

When using an M1 approach, we can expect worsened analyses outcomes when BEAMs are present, whether moderate or severe. Whilst M2 may cope with BEAMs innately due to the MV filtering process, the feature loss must be deemed acceptable. Nevertheless, neither method can acceptably handle BEAMs in data without some drawbacks. Thus, a novel approach may be necessary to prevent or at least, reduce the negative impact that BEAMs have on the data.

### Limitations and future research

Although we explained the rationale from an efficacy standpoint, we recognize the limitation of studying only one MVI (KNN) on the BEAMs phenomena. In reality, many MVIs exist, and their specific susceptibility to BEAMs were not investigated in this study. However, we expect that most MVI methods will produce similar outcomes anyway.

Additionally, the MV mechanisms and proportions were arbitrarily determined at 3 MNAR to 1 MCAR, which although not baseless, is not an absolute representation of real-world data. Thus, although our results did demonstrate the negative impact of BEAMs on MVI, we might expect different outcomes with real world data possessing different characteristics. Nonetheless, our intention was to point out that BEAMs can lead analysis astray.

Finally, performing MVI on data with BEAMs with either a global (M1) or batch-sensitized (M2) approach both results in poor outcomes. Hence, future research may be focused on developing a BEAMs-specific strategy that may ideally preserve the performance of M2 with the feature retention of M1.

## Conclusion

In summary, this study investigated the consequences of Batch-Effect Associated Missing Values (BEAMs) on data when performing Missing Value Imputation (MVI) using K-Nearest Neighbors (KNN) algorithm. The study found that the manifestation of BEAMs in data can lead to poor performance of MVI in both global (M1) and batch-sensitized (M2) approaches. It was also found that BEAMs lead to worsening performance in terms of either statistical analyses or the size of data, or both. Additionally, the results showed that batch effect correction alone does not eliminate the negative effects of BEAMs on MVI performance. The study highlights the need for the development of new strategies to handle BEAMs in data to improve the performance of downstream analysis.

## Materials and Methods

All methods were performed in the R programming language (version 4.2.2) and in the RStudio integrated development environment (version RStudio 2022.12.0+353), following appropriate guidelines and regulations.

### K-nearest neighbour (KNN)

There are various MVI methods that have been developed to accommodate different types of data. The simplest form of MVI is mean imputation, which replaces MVs with the mean of the corresponding feature. However, this method is not considered informative enough, so it is not used in this study. On the other end of the spectrum is the Random Forest (RF) algorithm(20,21), which is considered one of the most robust MVI techniques as it can perform well with almost all MV mechanisms(22). However, RF is computationally expensive and time-consuming because it requires numerous iterations.

A middle ground between mean imputation and RF MVI may be found with the K-nearest neighbor (KNN) algorithm. It is a simple machine learning method that replaces MVs with the mean of the K most similar data points(23). The nearest neighbors are typically determined by Euclidean distance, which is the distance between two points. Given two samples 1 and 2 measured on two variables x and y, this may be calculated by the formula:

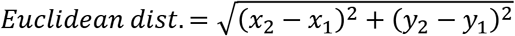

The KNN can be a good trade-off between the simplicity of mean imputation and the robustness of RF, while still being computationally efficient. KNN is not only fast and simple to perform, but it is also known for being one of the most effective MVI methods(24). An important parameter that influences the performance of KNN is the K value, which determines the number of nearest neighbors used for MV estimation. In our preliminary studies, it was shown that using the square root of the batch size for K was a conservative way to optimize self-batch imputation for retaining batch information to aid batch effect correction. In this study, the same principle was employed, and the square root of the batch size was used as the K value (✓10 = 3). The KNN algorithm from the “impute” R package, version 1.72.2 was used(25). By default, the algorithm performs mean imputation when the MVs exceed 50% in a row (in this case, feature). To assess pure KNN imputation, the threshold was set to 100% so that no mean imputation was introduced.

### Imputation strategies (M1 & M2)

In this study, two MVI strategies, M1 and M2, were applied. M1 is a typical global imputation strategy, where batch covariates are not considered. M2 is a batch-sensitized approach, in which MVI is performed within each batch. In a previous study, it was found that an M2 approach was generally superior as it improved BECA performance by retaining the batch covariate within the data for a more reliable correction(17). An MV threshold of 80% was used in this study, meaning that features with 80% or more MVs were removed from the data. This threshold was applied differently in M1 and M2 imputation strategies. With M1, this threshold was set across all batches. With M2, features must fulfill this threshold criterion in both batches to be retained in the data.

### ComBat

Many Batch Effect Correction Algorithms (BECAs) exist, such as ComBat(16), Batch Mean Centering(26), Harman(27), Surrogate Variable Analysis(28), each with different principles. ComBat is the most popular, ranking highly in several BECA evaluation studies(29–32). It uses an Empirical Bayes approach to estimate and standardize the means and variances between batches, effectively removing batch effects from the data(16). The implementation of ComBat used in this study was obtained from the “sva” R package, version 3.46.0(28).

### Data simulations and real-world data

To simulate data that may represent real-world data, we employed the method described in Shah et al. 2017 which used blockwise correlation and autoregressive correlation to create data containing features with correlations(33). The resulting dataset contained 20 samples with 900 features. For class and batch effects, we used the same method described in our previous study(17). Class effects were simulated for the first 60 features by a 1.5 times multiplication, and batch effects were simulated for the first 10 samples (known as batch 1) by addition and multiplication. This results in a dataset with two batches, each containing two classes. PCA was performed to visualize the artificial batch and class variances, before and after batch effect correction (Figure S1). To simulate the missingness found in real world data, MVs were then artificially inserted at a 3 MNAR to 1 MCAR ratio(11) at three different levels – (1) Control level: 30% MVs in both batches; (2) Moderate BEAMs: 20% MVs in one batch and 40% MVs in the other; and (3) Severe BEAMs: 10% MVs in one batch and 50% MVs in the other (Table 1). The MV simulations were performed 100 times.

**Table 1.**
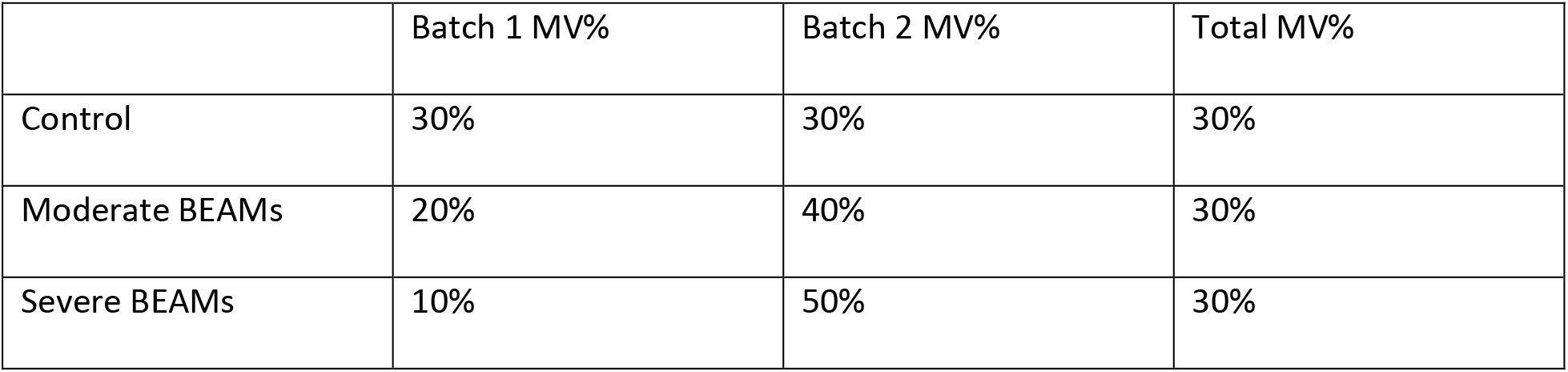
The three levels of batch MV proportions simulated.

|  | Batch 1 MV% | Batch 2 MV% | Total MV% |
| --- | --- | --- | --- |
| Control | 30% | 30% | 30% |
| Moderate BEAMs | 20% | 40% | 30% |
| Severe BEAMs | 10% | 50% | 30% |

This study used a real-world example, the Clinical Proteomic Tumor Analysis Consortium (CPTAC) study 6 benchmarking dataset, which consists of a yeast proteome as the background with human UPS proteins spiked in at different concentrations(18,19). We selected data from three machines at three spike-in concentrations to form a dataset with 3 batches, containing a total of 27 samples. The missing proportions are around 64%, 33% and 45% in the respective batches, amounting to 47% MVs overall (Figure 8). The data is available at: https://pdc.cancer.gov/pdc/browse?currentPath=%2525252FPhase_I_Data%2525252FStudy6&nonav=true.

### Evaluation methods

The following evaluation methods apply only to simulated data.

### Imputation accuracy

The accuracy of the MVI was assessed by the normalized root mean square error (NRMSE) metric, which compares the similarity of the imputed values to the true reference values. A lower NRMSE value suggests that the imputed values are more similar to the true values and therefore implies higher accuracy. The formula for calculating NRMSE is shown below, where x and y refer to the corresponding true and imputed values respectively, and sigma represents standard deviation.

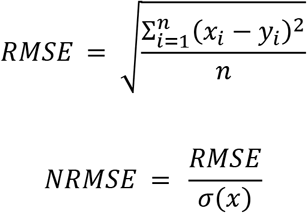

### Differential analysis

In the simulated data, class effects were introduced for the first 60 features. By performing a t-test on the initial data, the features that are truly differential can be determined and we may then identify the true positives (TP), false positives (FP), true negatives (TN) and false negatives (FN). The True Positive Rates (TPR) and False Positive Rates (FPR) can then be calculated from the imputed datasets. A higher TPR and lower FPR indicate better performance. TPR and FPR are calculated as follows:

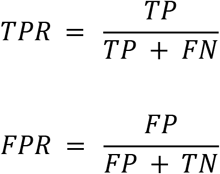

## Abbreviations

BE: Batch effect
BEAM: Batch effect associated missing value
BECA: Batch effect correction algorithm
CPTAC: Clinical Proteomic Tumor Analysis Consortium
FPR: False positive rate
KNN: K-nearest neighbors
LOD: Limit of detection
MAR: Missing at random
MCAR: Missing completely at random
MNAR: Missing not at random
MV: Missing value
MVI: Missing value imputation
NRMSE: Normalized root mean square error
PCA: Principal component analysis
TPR: True positive rate

## Data availability

The codes used to perform analyses in this study can be found in the GitHub repository: https://github.com/HarvardHui/BEAMs-paper.

The CPTAC study 6 dataset can be obtained from: https://pdc.cancer.gov/pdc/browse?currentPath=%2525252FPhase_I_Data%2525252FStudy6&nonav=true.

## Author contributions

HWHH performed analyses, developed figures and wrote the manuscript. WWBG conceptualized, supervised, provided critical feedback and wrote the manuscript.

## Acknowledgements

WWBG acknowledges support from an MOE Tier 1 award (RS08/21).

