## Supplemental Figures for "Uncovering the consequences of batch effect associated missing values in omics data analysis"

**Supplementary**

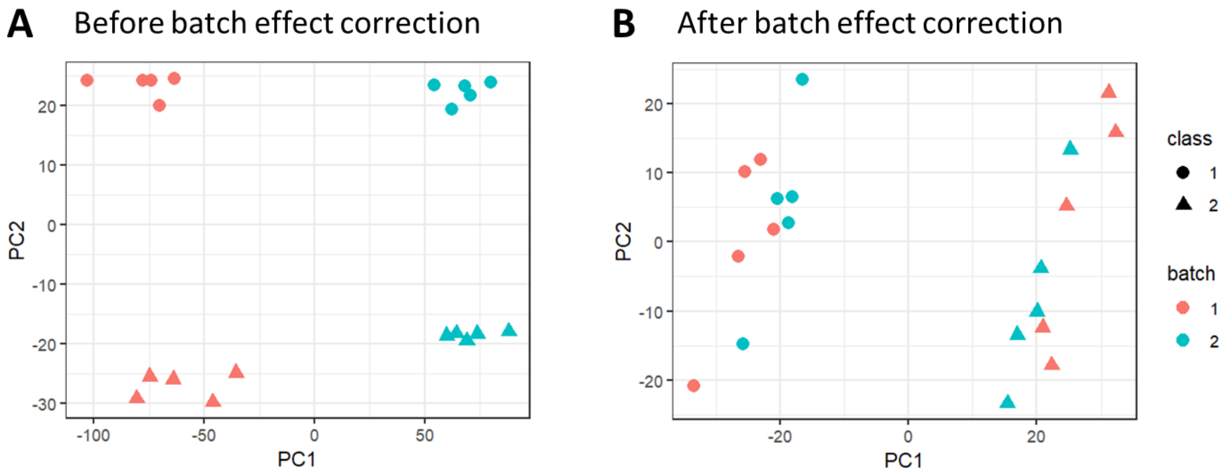


***Figure S1. Visualization of the simulated batch effect and class effect (A) before, and (B) after batch effect correction. Batch effect was visibly reduced after batch effect correction, whilst the class effect became the primary variance contributor.***

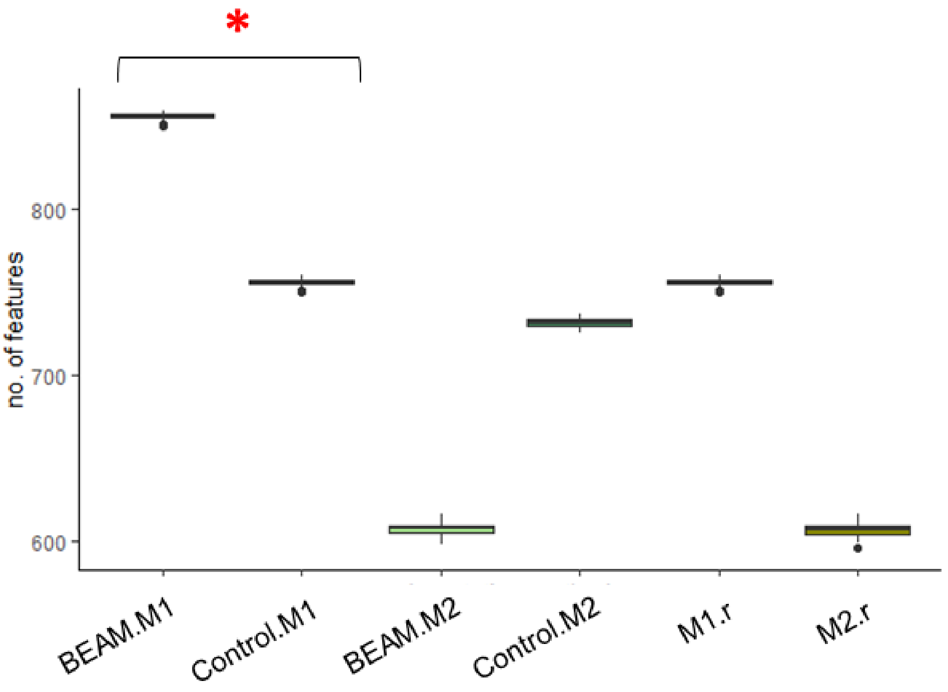


***Figure S2. Number of features retained after features with 80% or more MVs were removed in data with severe BEAMs (10:50% MVs). M1 and M2 with an "r" suffix refer to common features between BEAMs and Control within the same imputation strategy. Asterisks indicate significant difference ($p < 0.05$).***

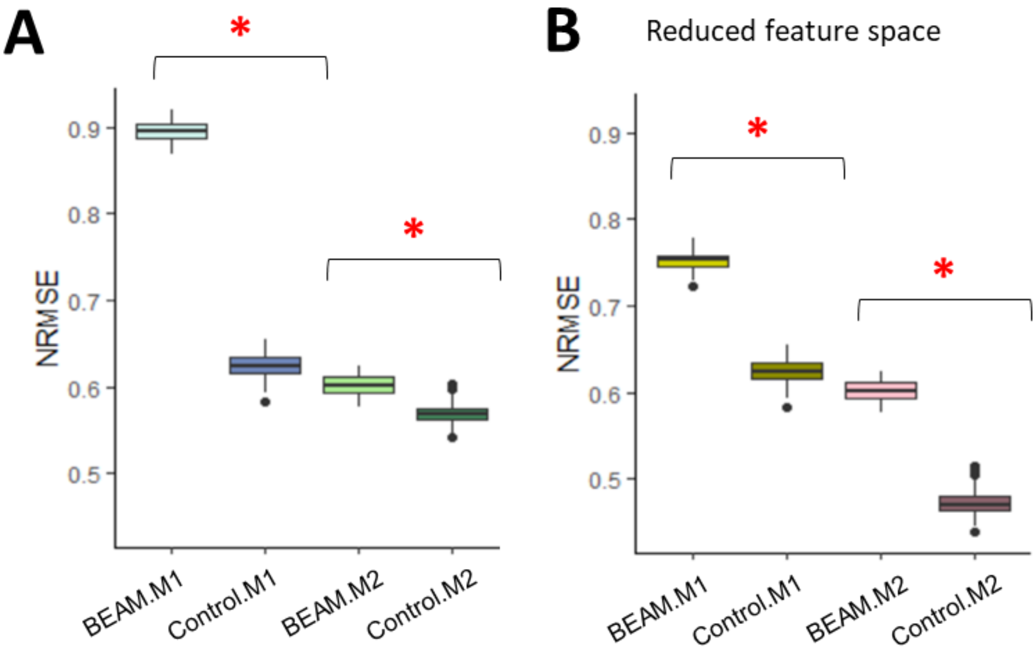


***Figure S3. NRMSE outcomes after MVI with and without severe BEAMs (10:50%). Asterisks indicate significant difference (p < 0.05). (A) Data are analyzed within their own feature space. (B) Data were reduced to the same features within the same imputation strategy (i.e., M1, M2).***

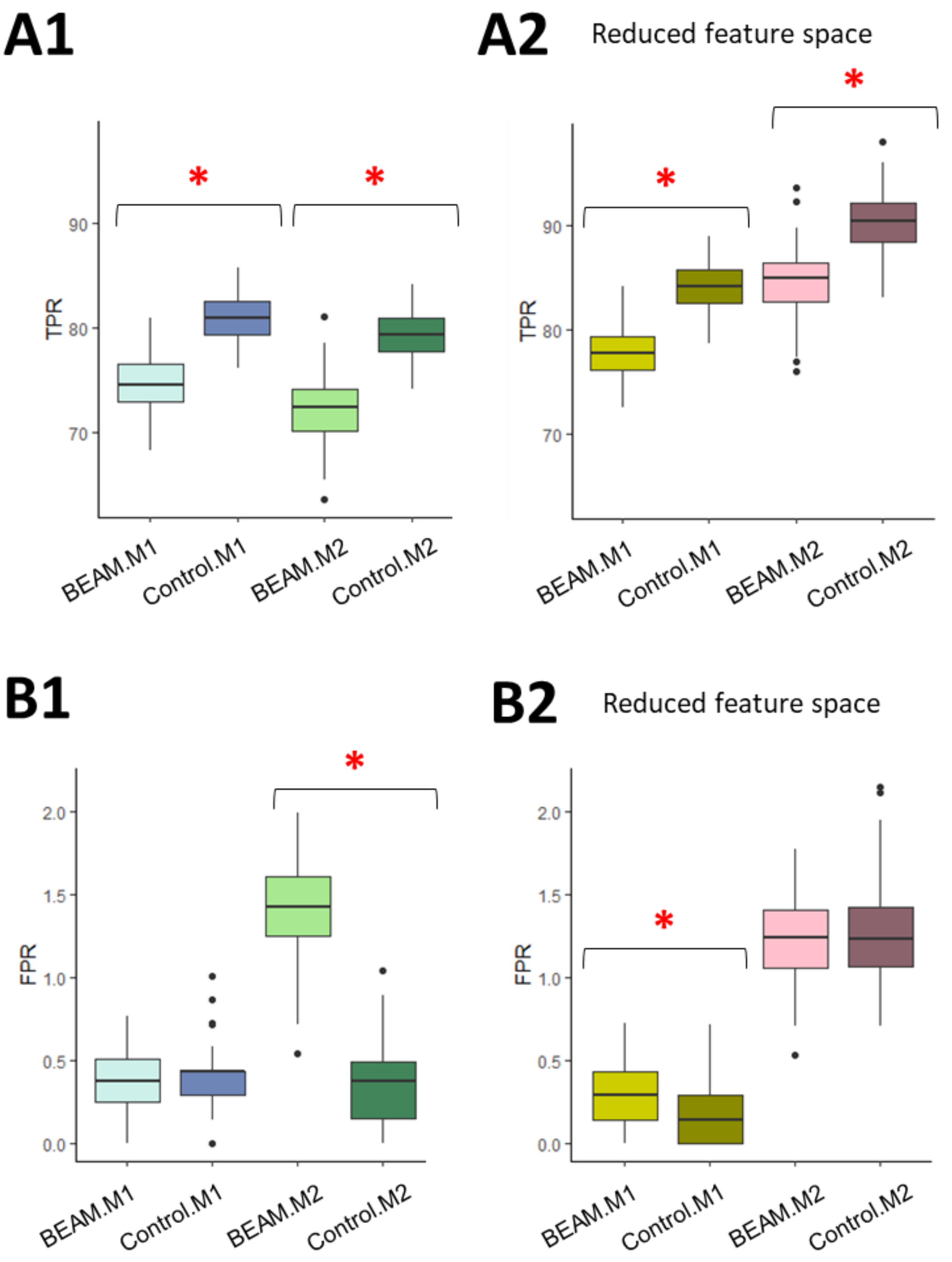


***Figure S4. (A) TPR and (B) FPR outcomes after MVI with and without severe BEAMs (10:50% MVs). Asterisks indicate significant difference (p < 0.05). (1) Data are analyzed within their own feature space. (2) Data were reduced to the same features within the same imputation strategy (i.e., M1, M2).***

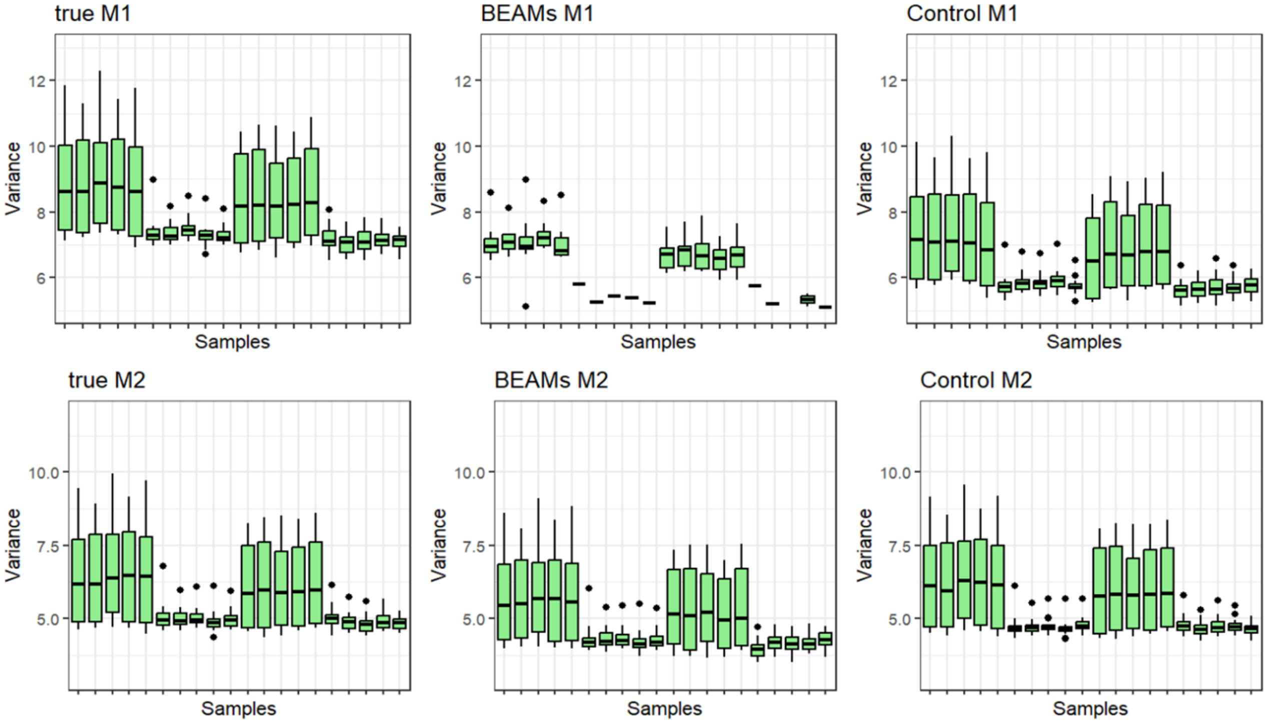


***Figure S5. Intra-sample variances of datasets from the final simulation iteration with moderate BEAMs (20:40% MVs). Data were reduced to the same features within the same imputation strategy (i.e., M1, M2).***
